# AXIS: Agentic Extraction and Integration of Simulation Scenarios for Quantitative Systems Pharmacology

**DOI:** 10.64898/2026.05.06.723244

**Authors:** Amirmohammad Kazemeini, Jose Prieto, Sanjana Balaji Kuttae, Anastasios Siokis, Chelsea Gallagher, Thomas Klabunde, Gurdeep Singh, Tommaso Andreani, Peyman Passban

## Abstract

Quantitative Systems Pharmacology (QSP) models are executable and mechanistic representations of biological systems that are increasingly used to design, evaluate, and help enrich clinical trials. Reusing a published model, however, requires knowing what each entity represents biologically (often inferable only from its relations to other entities) and what value it must take for a given scenario, information carried in the accompanying publication, not the model file. Agentic systems automate simulation setup from user-specified tasks over standardized parameter sets. In parallel, literature-mining tools extract reported quantities from publications. However, closing the loop, going from paper and model to an executable configuration that reproduces the published result, remains a manual curation task. Here, we present AXIS, a modular multi-agent framework that converts a published QSP model together with its companion publication into simulation-ready configurations. The framework integrates dependency-aware entity grounding, evidence-linked scenario extraction, semantic and structural entity search, sandboxed deterministic computation, and expert input on ambiguity. AXIS comprises three modules that ground the model’s entities, recover the scenarios described in the paper, and resolve each into concrete parameter assignments. We demonstrate the framework end-to-end on a published disease model, reconstructing the figure reported in its companion publication. AXIS recovers the reported scenarios, produces parameter assignments matching an expert-curated ground truth, and defers to an expert where a required quantity appears in neither source.

## 1 Introduction

Quantitative Systems Pharmacology (QSP) is a mechanistic modeling approach that quantitatively represents biological systems to understand disease processes and predict drug responses [1]. QSP has become a central discipline for modeling and simulation efforts aimed at de-risking clinical development programs and assessing the translational potential of novel drug assets. QSP models are typically formulated as systems of dynamical equations encoding causal assumptions about molecular pathways, cellular interactions, and physiological processes [2], and are commonly shared and archived using the Systems Biology Markup Language (SBML), a free, open, XML-based standard for representing model structure [3]. A published model therefore reaches a reader as two separate artifacts: an SBML file deposited in a public repository such as BioModels [4], and the publication that reports which simulations were run with it. The ability of QSP models to integrate diverse biological and pharmacological knowledge into mechanistic frameworks has made them an important component of model-informed drug development, supporting target evaluation, dose optimization, biomarker discovery, and translational decision-making across preclinical and clinical settings [5, 6].

Despite this value, regenerating a published QSP model’s reported simulation results from the information provided in the original study remains a significant challenge [7, 8], typically requiring substantial manual effort that complicates validating existing models, integrating them into new studies, or building upon prior work. While SBML effectively encodes model structure, it does not necessarily capture the simulation-specific settings needed to regenerate published results. Related standards such as the Simulation Experiment Description Markup Language (SED-ML) [9, 10] were introduced to describe simulation experiments in a software-independent way, including model changes, simulation procedures, and expected outputs. In practice, however, they are rarely authored for published models, as authors are not required to provide them, and the deposited files seldom carry the information needed to execute a scenario [11]. This gap gives rise to two key challenges.

The first challenge is the extraction of simulation scenarios. QSP models are rarely evaluated under a single experimental condition. Instead, the same model is executed under multiple conditions. Reconstructing these simulation scenarios requires manually identifying and extracting the corresponding settings from the accompanying publication, where they are often distributed across the text, figures, and tables, with each scenario requiring a distinct combination of entity assignments to reproduce the reported results. Entities may not be referenced directly, and even when they are, their values or units may differ from those encoded in the SBML file, requiring manual verification and unit conversion before the results can be reproduced. For example, a dose reported as 8 mg/kg every two weeks must reach the model as an absolute amount assigned to `DoseQ2W`, a parameter the publication never names and whose unit the model file does not declare [12].

The second challenge is establishing a semantic mapping between computational model entities and their biological meaning, without which downstream assignments fail silently. Model entities are expressed through complex chemical reactions and expressions in the model, and frequently lack sufficient semantic annotation, such that their biological interpretation cannot be determined directly from the model file [13]. An entity label alone rarely reveals its biological role (e.g., a production rate versus a degradation rate versus a transport constant); resolving this typically requires manual examination of the model such as its reactions and interconnections with other entities alongside information distributed across different sources, external databases, or the accompanying publication [14].

These two challenges extend beyond pharmacology: incomplete reporting of the information required to rerun a published simulation is a recognized barrier to reproducibility across computational modeling [7, 15].

Recent advances in large language model (LLM) agents have enabled the extraction of information from unstructured scientific literature using natural language, the integration of external tools and databases, and the execution of complex multi-step analytical workflows. Agentic systems such as Talk2Biomodels [16] and Talk2KnowledgeGraph [17] support natural language interaction with computational models and biomedical knowledge graphs, while Biomni-A1 [18], MEDEA [19] and BioMedAgent [20] combine reasoning with external tools and databases to support complex biomedical analyses, and MDCrow [21], NAMD-Agent [22] and DynaMate [23] automate molecular dynamics setup from user-specified tasks. The AI Scientist [24] further illustrates the potential of autonomous agents for iterative scientific reasoning and experimentation. These systems demonstrate the increasing applicability of LLM-driven agents to a broad range of biomedical research applications [25], yet none closes the loop from paper and model to executable configuration.

Building on these advances, we introduce AXIS, an agent-based framework for translating literature-derived simulation scenarios into reproducible, simulation-ready QSP configurations. The core technical contribution of AXIS is its specific decomposition-focused design. Rather than relying on ungrounded LLM calls to directly emit a complete model configuration, AXIS divides the task into a sequence of smaller, individually verifiable operations, each emitting an inspectable intermediate artifact, with the artifacts bound to one another. This design makes the pipeline grounded, traceable, and augmented by a domain expert at the point of ambiguity. Each stage is introduced here at the level of its inputs, outputs, and decision rule; the corresponding artifacts and worked examples are provided in the appendix.

## 2 Methodology

AXIS divides the top-level question of “How to reproduce the model results with scenarios mentioned in the accompanying paper” into three sub-questions, one per module: “How to construct a single, understandable model context with sufficient information (Semantic Profiler)”, “How to extract the scenarios of the accompanying paper relevant to the model of interest (Scenario Extractor)”, and “How to map individual free-text scenarios of the paper into the correct entities and values of the model (Scenario Mapper)”.

The pipeline’s sequential structure is also important: feeding the model context into scenario extraction ensures that only scenarios relevant to the model are selected, since a paper may describe multiple models. The mapping module likewise uses the model context together with one extracted scenario at a time, so that each scenario is mapped with full context of the model it concerns. An overview of the workflow is available in Figure 1.

**Figure 1.**
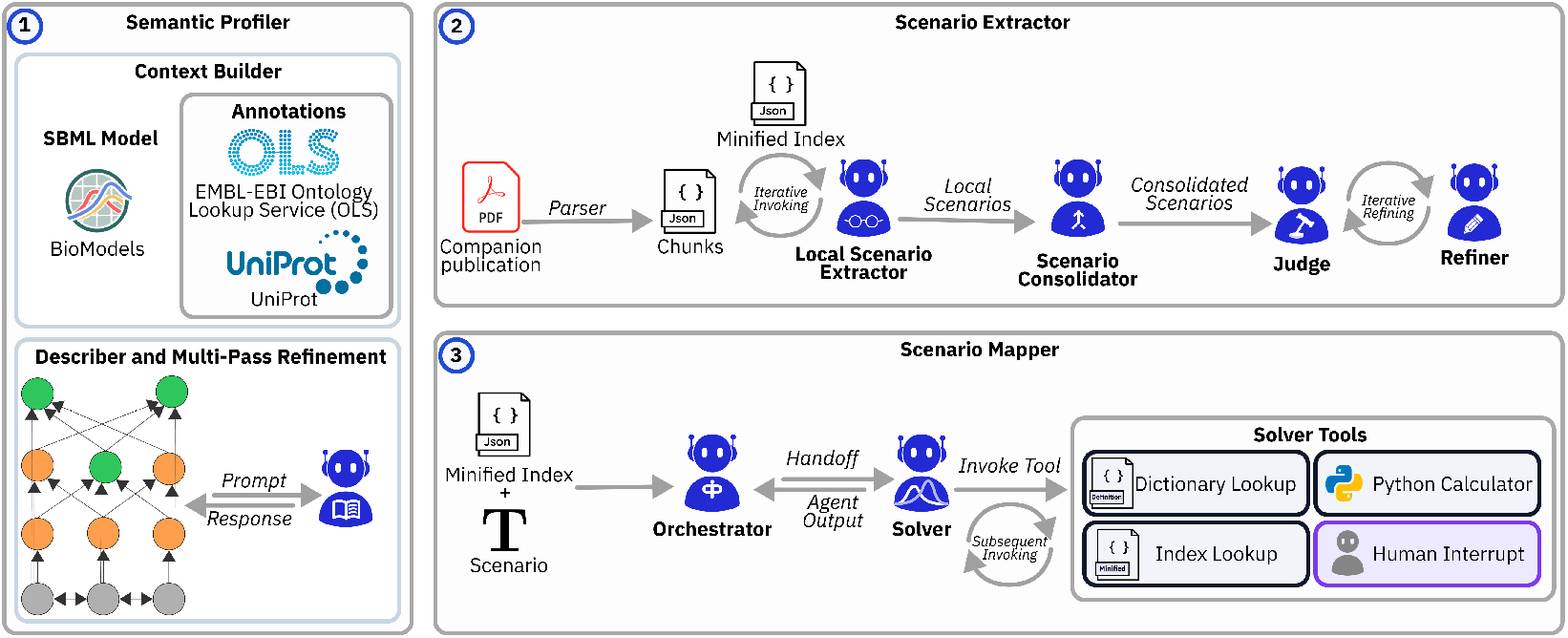
Overview of AXIS. The Semantic Profiler (1) grounds the model’s entities into an Entity Dictionary and its Minified Index over repeated refinement passes; node colour marks status: grey raw, orange refining, green complete. The Scenario Extractor (2) recovers the scenarios the companion publication describes for that model. The Scenario Mapper (3) resolves each into simulation-ready parameter changes.

As a running example, we use the Crohn’s disease IL-6 study of Dwivedi et al. [12] and its associated BioModels entry, BIOMD0000000537 [26], for the remainder of this paper, reproducing the result of the specific scenario shown in Figure 3a of that publication: “Treatment with 8 mg*/*kg anti-IL-6R*α* antibody administered intravenously every 2 weeks with a *K*_*d*_ of 1 nmol*/*L.” The reported readout is serum C-reactive protein (CRP) as a percentage of baseline.

### 2.1 Semantic Profiler

The first major module of AXIS is the Semantic Profiler, which takes a QSP model and generates biologically meaningful descriptions of its internal model entities. QSP models are typically stored as XML files, and many entity names are cryptic or difficult to interpret without additional context, such as `L` and `T` in BIOMD0000000913, or `C_free` in BIOMD0000000788. Even a seemingly descriptive name such as `kIL6RUnbind` in BIOMD0000000537 provides only partial information: the interacting species and direction, the compartment and reaction involved, and the initial value and unit are still missing.

To resolve this, the Semantic Profiler first augments each model entity with all its relevant context in a deterministic way (Mechanistic Context Builder), and then digests, interprets, and summarizes it in a semantically enriched and more human-readable JSON representation.

#### Mechanistic Context Builder

The goal of this component is to augment each model entity, that is, each parameter or species declared in the model, with the full relevant raw context generated as a structured JSON representation for next stage, LLM based interpretation. The information can be drawn from different sources such as the model XML file (for example, reactions and expressions if available) parsed via the BASICO [27] Python interface and external databases such as UniProt [28] and the Ontology Lookup Service (OLS) [29] for entity description, or context from the paper. We focus on the model and external databases for this paper. For the external databases, we expanded upon the annotation retrieval tool introduced by Talk2Biomodels [16].

In the XML model, for example, the entity “IL6{serum}” is paired with UniProt ID P05231. Retrieving that annotation adds the name “Interleukin-6”, which provides a better biological interpretation. IL6{serum} is also involved in several reactions, each of which references a function ID for its rate formula that must be decoded to retrieve a readable form. Further explanation and example output of the Context Builder are available in Appendix A.1.

By fetching this verified context prior to inference, the Context Builder grounds the LLM in established biological records rather than leaving it to speculate about the meaning of an entity. Nasri et al. [30] report the same effect in a different domain: deterministic access to authoritative records, rather than model recall, is what makes agentic scientific workflows robust. The direct result of this stage is an intermediate Enriched Mapping artifact, which cleanly aggregates the governing reaction equations, passthrough initial values, and unified annotations to serve as the foundational context for the downstream Describer.

#### Describer and Multi-Pass Refinement

Even though the output of the Context Builder is represented in a structured JSON format, it remains a long, raw technical description of the model rather than a clear biological explanation. For example, it records only that parameter `kIL6RUnbind` appears in the reaction “Ab{serum} + sR{serum} = Ab_sR{serum}”.

The describer uses an LLM to analyze the context produced by the Context Builder. During this process, the LLM is supplied with the full extracted context of an entity, in which some of the unprocessed raw context, such as reactions and annotations, is transformed into new fields, such as `biological_identity` and `definition` and role in the model, while existing entries may be augmented with further explanatory information (see Appendix A.2 for details).

Since biological models are highly interconnected, the interpretation of one entity may depend on the meaning of several other entities. Consequently, an entity definition generated by the describer may remain incomplete when it refers to model symbols that have not yet been biologically resolved, which triggers the refinement pass. For example, the initial definition of `kIL6RUnbind` reads “kIL6RUnbind controls the dissociation of Ab_sR into Ab and sR” without disambiguating the entities it names.

In each refinement pass, the Describer re-issues the original raw context, now augmented with a “Context from Related Entities” block. To keep the prompt compact, this block carries only the latest biological identity and definition of the missing dependencies, pulled from a running cache updated after every pass. The refinement loop terminates when all entity dependencies are resolved, when no progress is observed, or when the `max_passes` threshold is reached. For example, the definition of `kIL6RUnbind` after resolving dependencies may be: “kIL6RUnbind is the kinetic parameter controlling the dissociation of the antibody-soluble receptor complex Ab_sR into free antibody Ab and soluble receptor sR.”

Refinement serves not merely to rewrite definitions in more fluent language but to ensure each entity’s description is complete and understandable on its own, without consulting other entities. See Appendix A.3 for the full example.

The Semantic Profiler emits two artifacts: the *Entity Dictionary*, which holds one full definition per model entity, and the *Minified Index*, a compressed one-line description of each entity derived from it. The two artifacts therefore serve complementary purposes. The Entity Dictionary supports exact lookup and comprehensive inspection of a known model entity, whereas the Minified Index supports efficient semantic grounding and candidate matching between natural-language statements in the publication and cryptic model entity names. Appendix A.4 gives both artifacts for the same entity.

### 2.2 Scenario Extractor

Scenarios are the representation that connects experimental descriptions in the literature to executable configurations of the corresponding QSP model. The Scenario Extractor reads the companion paper and identifies experimental or simulation settings described in the text. Its input is the paper text together with the Minified Index, and its output is a structured collection of candidate scenarios. In the current implementation, extraction is performed only from textual content; figures, diagrams, and other visual elements are not yet included. The Scenario Extractor follows a three-block architecture: local extraction, global consolidation, and iterative judge-refine loop.

#### Local Scenario Extraction

The paper is divided into small, ordered chunks of approximately 320 tokens. Candidate scenarios are evaluated using overlapping windows composed of two consecutive chunks, with an overlap of one chunk between adjacent windows. This design avoids presenting the entire paper to the LLM in a single prompt and allows the extractor to detect locally described experimental or simulation conditions while preserving limited surrounding context. For each window, the extractor evaluates whether the text contains a scenario that could be instantiated and executed in the QSP model. Relevant information may include the intervention, administered dose, route of administration, and similar execution settings.

#### Global Scenario Consolidation

Because local windows overlap and related information may be expressed differently across the paper, the initial extraction stage may produce duplicate, incomplete, or partially overlapping scenario candidates. The global consolidation module compares the locally extracted candidates, merges equivalent descriptions, removes redundancy, and constructs a coherent scenario set. For example, “8 mg/kg anti-IL-6R every 4 weeks” and “anti-IL-6R*α* antibody administered Q4W” may refer to the same treatment scenario. The consolidation module determines whether these descriptions are semantically equivalent and, when supported by the source text, merges them into a single scenario representation. The module may also identify candidates that are too broad to represent a single executable scenario, such as “dose-response experiment”. This description may correspond to several distinct dose levels or parameter sweeps. In such cases, the candidate is decomposed into separate scenarios when the paper provides sufficient evidence to identify the individual experimental conditions.

#### Scenario Judge and Refiner

After global consolidation, the Judge evaluates the complete scenario set for remaining inconsistencies or omissions. These include duplicated scenarios, unsupported details, missing experimental conditions, improper merging, overly broad descriptions, or scenarios that should be divided into multiple executable configurations. Rather than regenerating the complete scenario set, the Judge requests corrections that the Refiner applies to the affected entries. This constrained correction strategy improves traceability and reduces the risk that previously supported information will be altered or removed during revision. This loop is bounded to three iterations. The objective is not to repeatedly rewrite the scenarios for stylistic improvement, but to produce a concise and consistent scenario set. Each scenario is assigned an ID together with the IDs of the relevant paper chunks, which preserves traceability throughout the process; Appendix A.5 shows the stored form.

### 2.3 Scenario Mapper

Once the scenarios have been extracted, they are passed to the Scenario Mapper. The mapper takes three inputs: a free-text scenario, the Minified Index, and the Entity Dictionary. It transforms these inputs into a simulation-ready representation containing the concrete parameter and species changes required to execute the scenario in the QSP model.

To perform this conversion, the mapper identifies the relevant model entities and assigns the values, units, and execution settings needed to reproduce the reported experiment or simulation. For example, a paper may report a weight-normalized dose (such as 8 mg/kg) whereas the model expects a total administered amount (such as `Dose` = 560 in mg).

Because this process may require entity identification, semantic matching, unit conversion, dependency resolution, and management of unresolved information, the mapper is implemented using two key cooperating agents: the Orchestrator agent and the Solver agent.

#### Orchestrator agent

The Orchestrator agent determines which model-level changes are required to represent the extracted scenario. It reads the scenario together with the Minified Index and identifies the model entities that are potentially relevant. It then decomposes the mapping problem into a set of discrete work orders.

Work orders are dispatched individually to reduce conflicts between simultaneous modifications and to make each mapping decision independently traceable. This design also supports human augmentation when a specific task cannot be resolved with the given context.

For the scenario “Treatment with 8 mg/kg anti-IL-6R*α* antibody every two weeks with a *K*_*d*_ of 1 nmol/L”, the Orchestrator emits two work orders: one targeting `DoseQ2W` for the unit conversion, and one targeting `kIL6RUnbind` for the affinity constraint. The Orchestrator therefore focuses on determining *what* must be resolved or changed based on the given context and it does not perform the underlying calculation or decide how the transformation should be executed.

#### Solver agent

Once the required tasks have been identified, the Solver agent determines how each work order should be completed. The solver has access to a set of tools that support entity retrieval, semantic search, deterministic calculation, and expert intervention.

**Dictionary Lookup** retrieves an entity’s complete semantic and numerical profile from the Entity Dictionary by exact name. **Index Lookup** resolves an entity name from a concept described in the scenario by matching it against the Minified Index; this simpler search is served by a smaller model, GPT-4.1-nano. **Human Augmentation Interrupt** is raised with a proposal when a mapping decision cannot be resolved safely from the paper and model alone, for example when an entity’s unit is missing. **Python Calculation Tool** performs deterministic numerical operations in a sandboxed environment, particularly unit conversion and arithmetic transformations.

The complete mapping process may involve several sequential tool calls. A typical sequence includes Index Lookup to identify candidate entities, Dictionary Lookup to inspect the selected entity, the Python Calculation Tool to derive the required value, and, when necessary, a Human Augmentation Interrupt to resolve missing information. Appendix A.6 provides an end-to-end example showing how a scenario is processed by the mapper and how these tools are invoked sequentially to produce a simulation-ready representation.

## 3 Results

Evaluating a system of this kind is challenging because no established gold-standard dataset exists for literature-to-QSP scenario extraction, and directly comparable methods are limited. We therefore adopted an expert-replication strategy. Domain experts first performed the extraction and mapping process manually, following their standard workflow for interpreting publications and reproducing QSP simulations. We then applied AXIS to the same model–publication pairs and compared its outputs with the expert-curated artifacts. Success here means being able to regenerate the scenario result from a starting point of having raw paper and model. Hence, we demonstrate it by executing the scenarios behind Figure 3a of the BIOMD0000000537 paper [12, 26] on Crohn’s disease.

To start, the Semantic Profiler processes the model and generates the semantic artifacts. Afterwards, the Scenario Extractor receives the paper PDF and the semantic artifact to find the relevant scenarios. Then we select the three scenarios corresponding to Figure 3a from the Scenario Extractor output and pass them to the Mapper unchanged. The final stage is executing the model with those scenarios to compare the simulated output with Figure 3a.

We walk through one example mapping step by step. The paper reports 8 mg/kg dosing every 2 and every 4 weeks, with a *K*_*d*_ of 1 nmol/L, compared against placebo. The Scenario Extractor creates three scenarios of “Treatment with 8 mg/kg anti-IL-6R*α* antibody administered intravenously every 2 weeks with *K*_*d*_ of 1 nmol/L” and “Treatment with 8 mg/kg anti-IL-6R*α* antibody administered intravenously every 4 weeks with *K*_*d*_ of 1 nmol/L” and “placebo treatment”, decoupling the merged description into separate full scenarios.

For the every-2-weeks scenario, the Orchestrator generates two work orders of `DoseQ2W` (mg) and `kIL6RUnbind` (1/h) for the Solver agent. The `DoseQ2W` work order requires the Solver agent to verify the proposed value for the missing body weight and dose unit from the user, after which it converts the dose to 560. To resolve the `kIL6RUnbind` work order, the relation *k*_off_ = *K*_*d*_ · *k*_on_ is computed by retrieving the missing association rate constant *k*_on_, which the model stores as kIL6RBind = 1000, and combining it with the literature-reported *K*_*d*_, giving a value of 1000. This illustrates the Solver’s ability to chain Dictionary Lookup, Index Lookup, and the Python Calculation Tool to resolve parameters that are coupled through reaction kinetics.

The remaining two scenarios resolve the same way. For the every-4-weeks dosing, the only difference is the target dose parameter, which is `Dose`, and the rest of the process is the same. Placebo treatment does not require any parameter change, which AXIS also successfully handles.

The parameter assignments match the expert-curated ground truth, and executing them reproduces the published curves (Figure 2). A small non-reasoning LLM (GPT-4.1-mini) suffices for this task.

**Figure 2.**
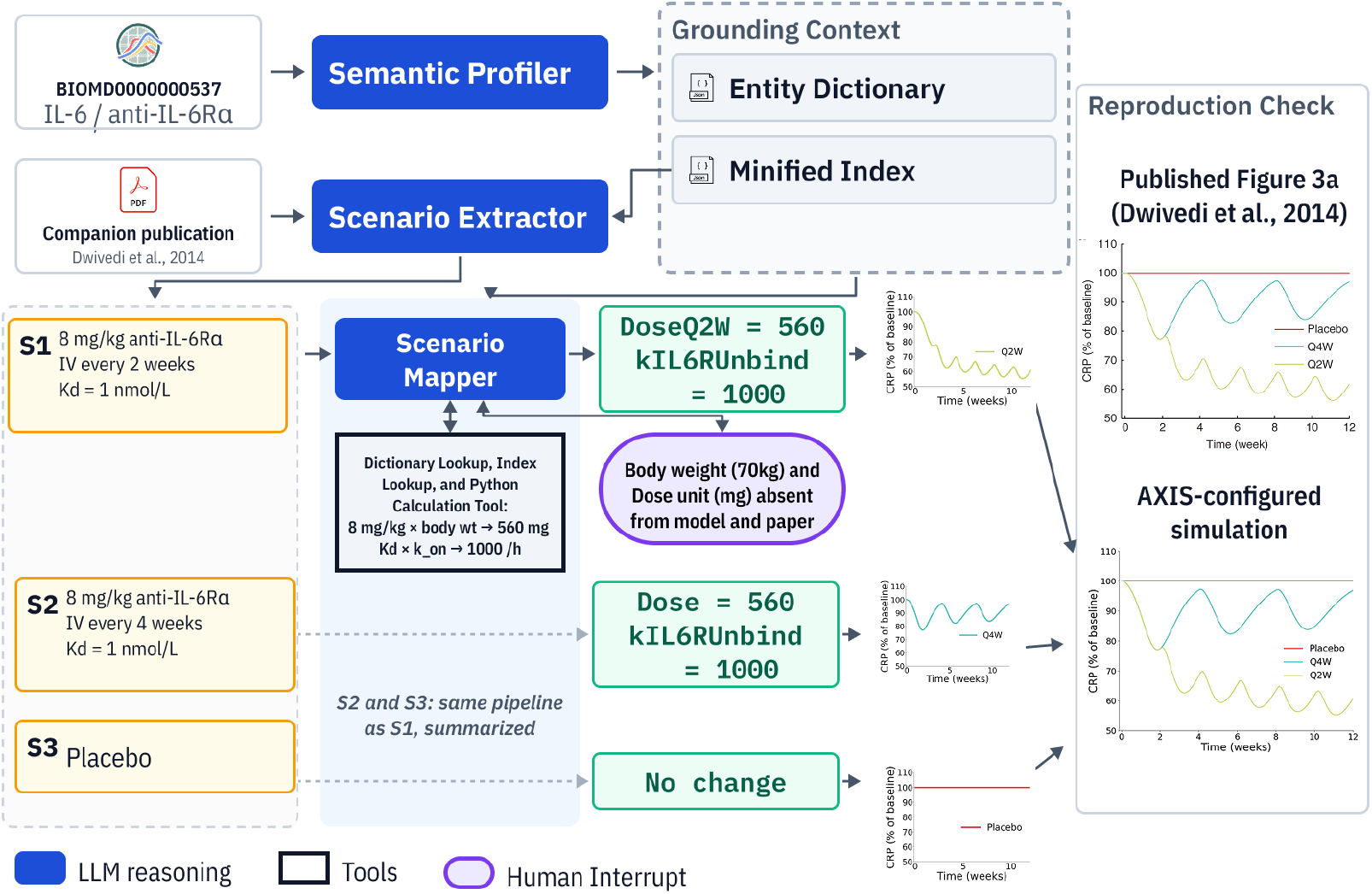
End-to-end reproduction of Figure 3a of Dwivedi et al. [12]. Configurations recovered by AXIS from the BIOMD0000000537 SBML file and its publication, executed for the three published arms: every two weeks (Q2W, S1), every four weeks (Q4W, S2), and placebo (S3). Body weight and the DoseQ2W unit, absent from both sources, are requested from the expert (purple) with a standard value proposed as default. S1 is drawn in full as a worked example; S2 and S3 resolve the same way and are drawn in summary.

Our framework can be integrated into existing model-simulation assistants such as Talk2Biomodels [16], supplying a grounded configuration before simulation and opening new directions for literature-grounded model reuse. To isolate the difference, we regenerate the every-two-weeks result with Talk2Biomodels. As it has no Scenario Extractor, we pass it the same pre-Mapper scenarios with the same backing LLM, restricting the comparison to the mapping step.

As shown in Table 1, Talk2Biomodels with the same backing model could not reach the correct configuration due to missing unit conversion and incorrect parameter extraction. Upgrading the backing LLM to a reasoning model recovers parameter identification in most runs, since the name is close to self-explanatory, but the dose mapping still fails in half the runs because the information needed to ground it is absent from both sources. To isolate this, we tested BIOMD0000000913 [31], whose entity names are not self-explanatory. The scenario “Treatment with a stabilized combination chemotherapy formulation that exhibits half the baseline degradation rate” requires setting `gamma` to 0.45. Neither backing model reaches it. Both halve a different parameter instead, most often `d`, the maximum *tumor* clearance rate rather than the *drug*’s, and in every run the submitted value is exactly half that parameter’s initial value. The instruction and the arithmetic are therefore recovered throughout; only the binding to a model entity fails. See Appendix A.7 for the full run logs.

**Table 1.** Scenario resolution across systems. A configuration is counted as correct when every required entity is named as it appears in the model file, with the correct value, and no other entity is changed. All Talk2Biomodels runs are supplied with the scenario extracted by AXIS. No run uses a seed, and all GPT-4.1-mini runs use temperature 0; o3 exposes no temperature parameter. Each count reports how many of the ten runs show that property, so a run may be counted more than once.

| System | Backing LLM | Outcome across 10 runs |
| --- | --- | --- |
| *BIOMD0000000537, every-two-weeks dosing (DoseQ2W = 560, kIL6RUnbind = 1000)* |  |  |
| AXIS | GPT-4.1-mini | correct configuration (10/10); identical across runs |
| Talk2Biomodels | GPT-4.1-mini | correct configuration (0/10); no dose unit conversion (10/10); kIL6RUnbind not resolved (10/10); dose written to Ab (2/10); error: entity not found (5/10) |
| Talk2Biomodels | o3 | correct configuration (4/10); kIL6RUnbind resolved (9/10), missed (1/10); both Dose and DoseQ2W changed (2/10); inconsistent incorrect dose value, such as 4000, 480, 1296 (3/10) |
| *BIOMD0000000913, stabilized chemotherapy (gamma = 0.45)* |  |  |
| AXIS | GPT-4.1-mini | correct configuration (10/10); identical across runs |
| Talk2Biomodels | GPT-4.1-mini | correct configuration (0/10); gamma never reached (10/10); halved d instead (10/10) |
| Talk2Biomodels | o3 | correct configuration (0/10); gamma never reached (10/10); halved d instead (6/10), s, u, beta, w (1/10 each) |

## 4 Discussion

We presented AXIS, an agentic framework for converting simulation scenarios described in QSP publications into structured, simulation-ready model configurations. AXIS combines semantic profiling of SBML models, dependency-aware entity refinement, scenario extraction, and grounded agentic mapping supported by deterministic calculations and human augmentation. By decomposing the workflow into grounded and traceable stages, the framework reduces unsupported reasoning and makes intermediate decisions available for expert inspection.

The current implementation nonetheless has several limitations. It assumes that models are available in a BASICO-compatible format such as SBML, whereas many industrial QSP workflows rely on custom code, proprietary platforms, or alternative simulation environments. The Scenario Extractor is limited to textual content, which excludes visual aspects of the paper as well as supplementary material. The performance of the Mapper depends on both the complexity of the scenario and the underlying LLM. Establishing generalizability and robustness across therapeutic areas, model structures, publication styles, and simulation platforms will require evaluation and refinement on a comprehensive dataset of diverse QSP papers. Future work will address each of these: additional model formats and execution engines; figures, tables, supplementary materials, and existing SED-ML specifications; and a curated expert-generated dataset built with QSP specialists, which would also enable benchmarking across backing models. We also plan to strengthen scenario disambiguation and to validate generated configurations automatically through model execution.

Automating the path from publication to executable configuration also carries a risk that this work documents directly: an incorrect configuration executes without error and produces a plausible trajectory, so a mapping error can propagate silently into downstream analysis. Where a scenario is ambiguous or under-specified in both sources, AXIS therefore defers rather than imputing a value it cannot ground, supporting expert review of model reuse rather than replacing it. Each of the three modules can be deployed on its own, composed end to end, or run alongside existing assistants such as Talk2Biomodels. We invite others to extend the framework, and to help assemble the corpus of paper and model pairs this task will need.

## Acknowledgments

We acknowledge Julia Deichmann for her in-depth discussions and help with the IBD proprietary model from Sanofi.

## Funding

This study was funded by Sanofi.

## Conflict of Interests

AK, JP, AS, CG, TK, TA and PP are Sanofi employees and may hold shares and/or stock options in the company.

## A Method Examples and Supporting Artifacts

Appendices A.1–A.4 walk through the artifacts produced by the Semantic Profiler, from the Context Builder’s structured model output (A.1) and its enrichment by the Describer (A.2), through the refinement of an under-specified entity (A.3), to the resulting Entity Dictionary and Minified Index (A.4). Appendix A.5 gives an example of an extracted scenario, and Appendix A.6 traces its complete mapping into simulation-ready parameters. Finally, the Talk2Biomodels baseline run logs referenced in Section 3 are given in Appendix A.7.

### A.1 Structured Model Output from the Context Builder

The Context Builder loads the model through the BASICO [27] interface and reads every model variable, both parameters and species, into a single record set. The two are read from separate model tables and carry different fields. A species is read with its initial concentration, its declared unit, and the compartment in which it is declared, and is keyed by its display name: the bare species name when that name is unique, and `name{compartment}` when the same name is declared in more than one compartment, so a species present in several compartments is carried as a separate record for each, with its own initial value, unit, and reaction set. A parameter is read with its initial value and declared unit and is keyed by its bare name, such as `Dose` or `kIL6RBind`, because parameters are model-level values that carry no compartment attribute.

Every entity of either kind is then matched against every reaction in the model by word-boundary matching on the reaction scheme, the species mapping, the reaction name, the rate function, and the reaction identifiers, and each matching reaction is attached to that entity’s record. The same matching is applied to assignment and rate rule expressions and to event triggers and assignments, so entities in models whose dynamics are expressed as rules rather than reaction kinetics are covered as well. A species additionally carries its own rate rule expression where one is defined, and a parameter additionally carries a reverse index of every expression elsewhere in the model that mentions it, which supplies context for a parameter that appears in no reaction scheme of its own. An entity that is purely reaction driven simply ends up with an empty rule context.

Annotations are retrieved for species only, from the MIRIAM (Minimum Information Required in the Annotation of Models) annotations recorded in the model file, which are returned as identifiers.org links. Each link is split into a database name and a database-specific identifier, grouped by database, and resolved to a human-readable label: UniProt [28] for protein entries, and the Ontology Lookup Service [29] for the remaining ontologies. Parameters carry no such annotation, so a parameter’s context consists entirely of the reactions and expressions in which it appears and of the entities named there. Units are carried for both kinds with their provenance, recording whether a unit was read from the model or is absent from it, alongside the model’s own declared unit system, since that unit system is not fixed across BioModels. This lets a downstream consumer refuse to compute with a unit that was never declared rather than infer one.

The example below is the assembled *Enriched Mapping* produced by the Context Builder. BASICO supplies the individual model tables and the raw annotation links; the reaction attribution, the rule and event context, the resolved annotation labels, and the per-entity aggregation are produced by this stage.

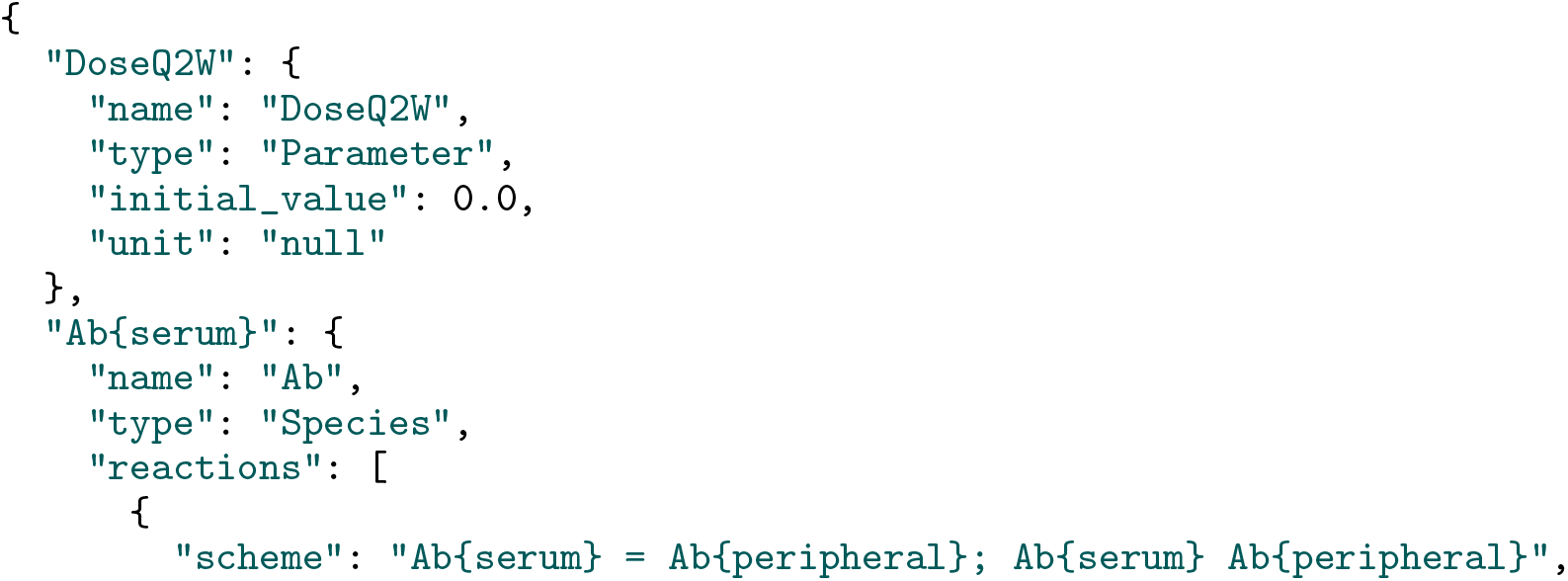
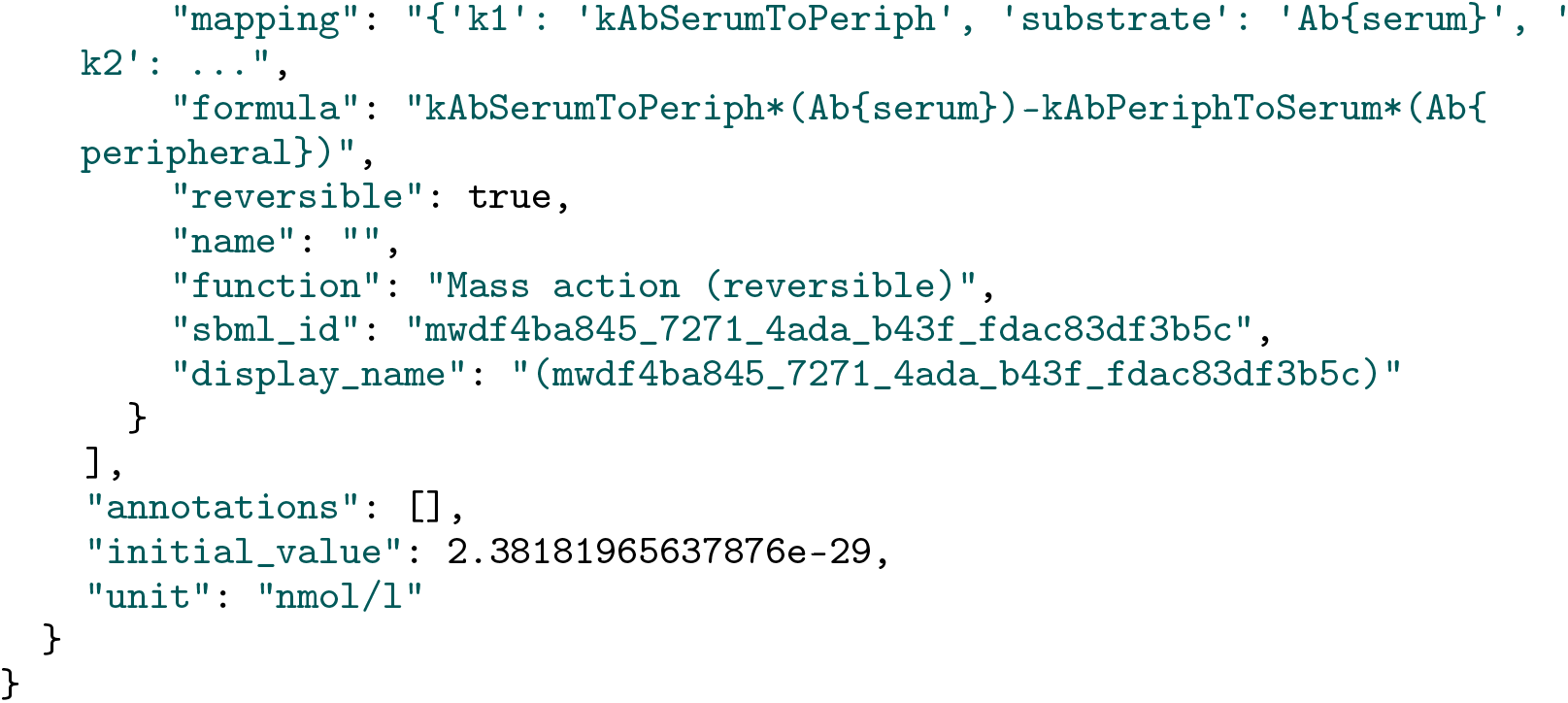

### A.2 JSON Representations Before and After the Describer

The mechanistic Context Builder may produce the following representation for kIL6RUnbind:

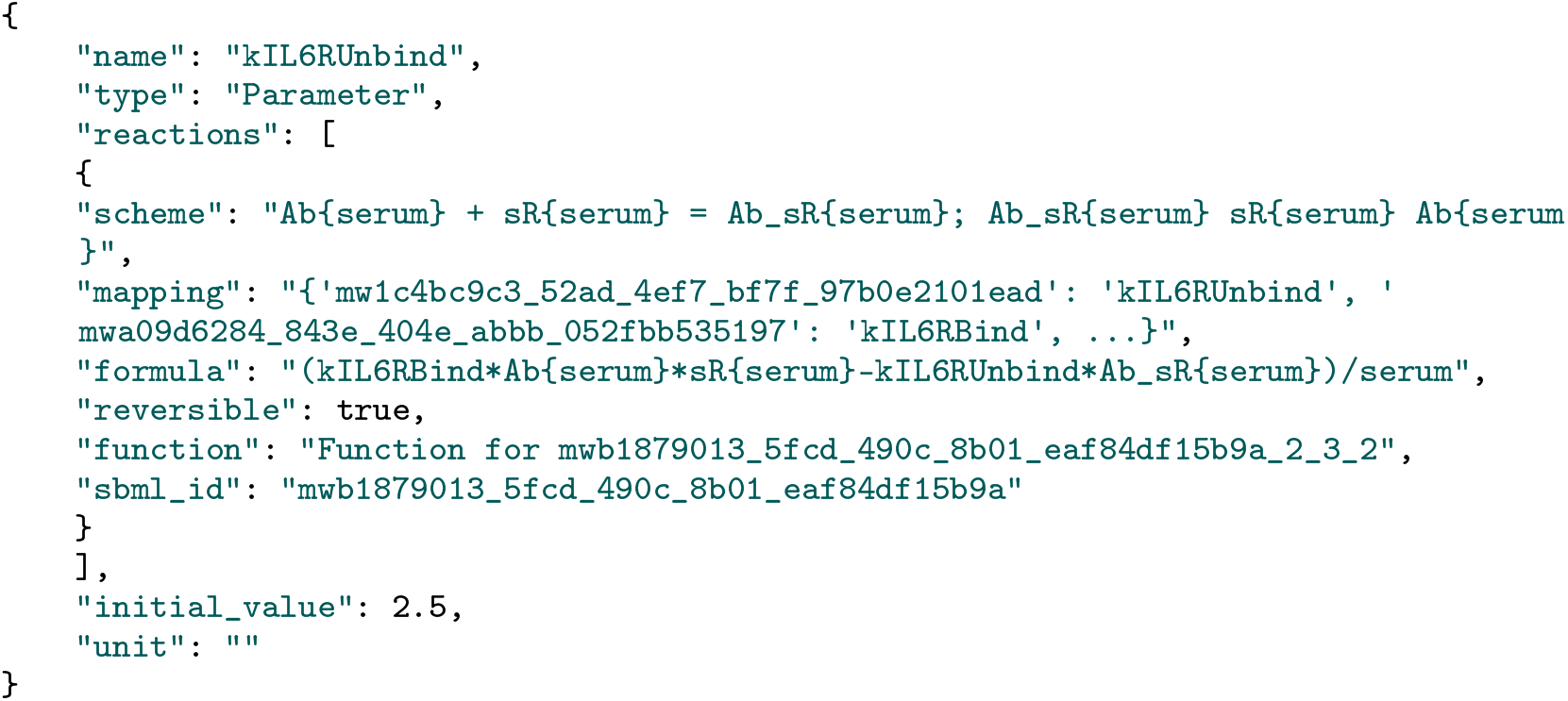

Although this representation contains useful mechanistic information, much of it remains difficult for a human reader to interpret. After processing this representation, the entity describer produces a semantically enriched JSON object such as the following:

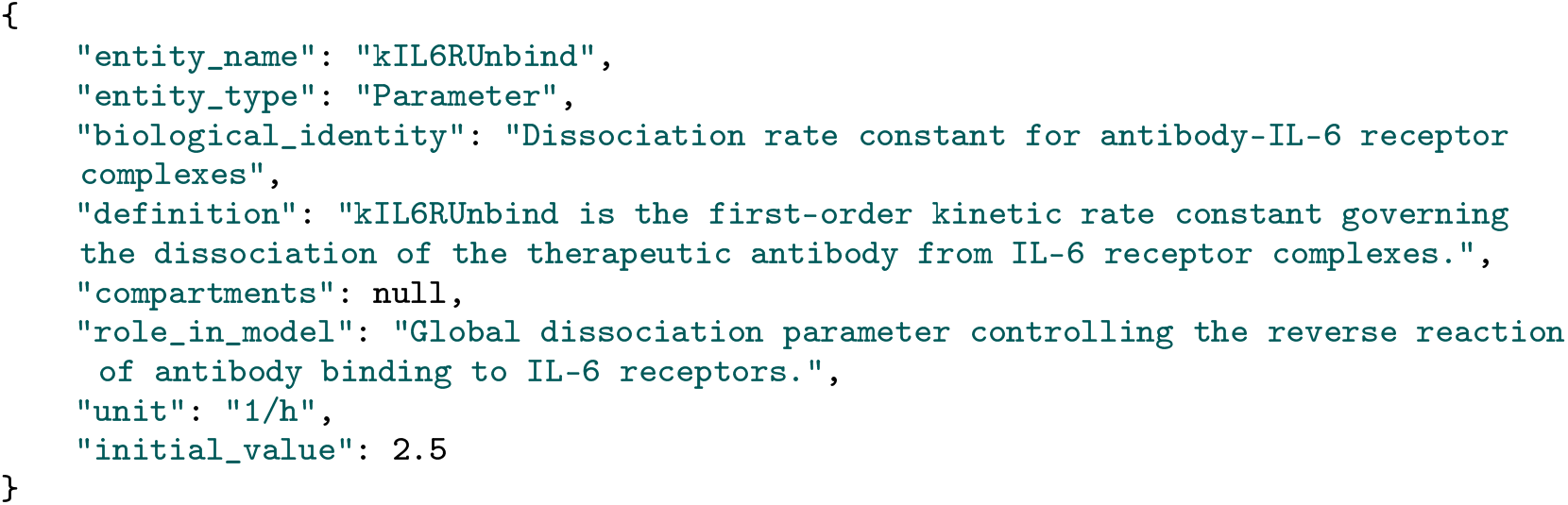

The enriched representation retains the original numerical and structural information while replacing cryptic identifiers with fields that explicitly describe the entity’s biological identity, mechanistic definition, model role, and associated compartments. This output is easier for domain experts to inspect and can also be used by downstream agents for natural-language scenario mapping.

### A.3 Refining an Under-Specified Entity

Some entities cannot be fully described in a single pass because their preliminary definitions reference other model symbols whose biological meaning the Semantic Profiler has not yet resolved. This subsection illustrates how such an under-specified entity is detected and subsequently refined, using the parameter `kIL6RUnbind` as an example. The Semantic Profiler may identify that this parameter occurs in the following reaction:

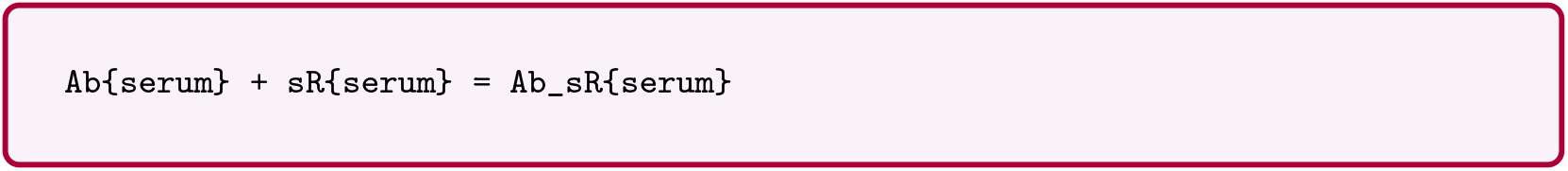

However, the biological meanings of `Ab{serum}`, `sR{serum}`, and `Ab_sR{serum}` remain unresolved, so they constitute unresolved dependencies of the definition of `kIL6RUnbind`. A definition written from this reaction alone is insufficient for both human reviewers and down-stream scenario-mapping agents, because the biological entities involved remain unclear. The Describer therefore marks the entity NEEDS_REFINEMENT and records the unresolved symbols in an `unresolved_dependencies` field.

The following intermediate representation shows an entity that has been partially interpreted by the Describer but still depends on unresolved model symbols:

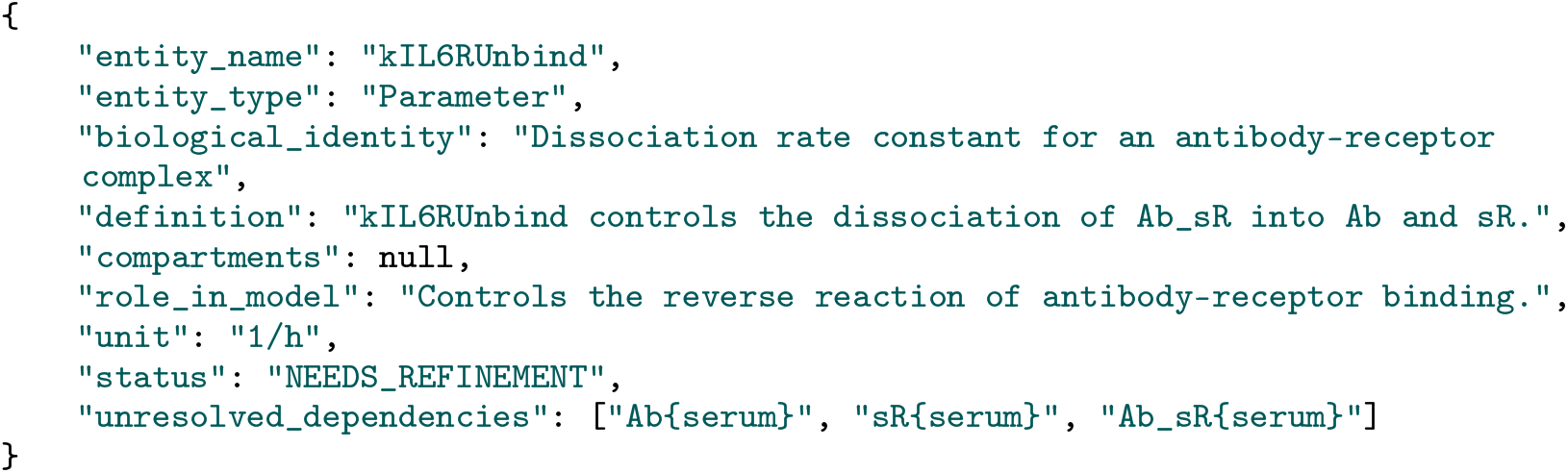

During the subsequent refinement pass, the Describer does not receive only the name and preliminary definition of `kIL6RUnbind`. It also receives the available semantic descriptions of every entity listed under `unresolved_dependencies`, here `Ab{serum}`, `sR{serum}`, and `Ab_sR{serum}`. With this added context the Describer rewrites the definition and names the biological entities the reaction involves, giving the resolved representation:

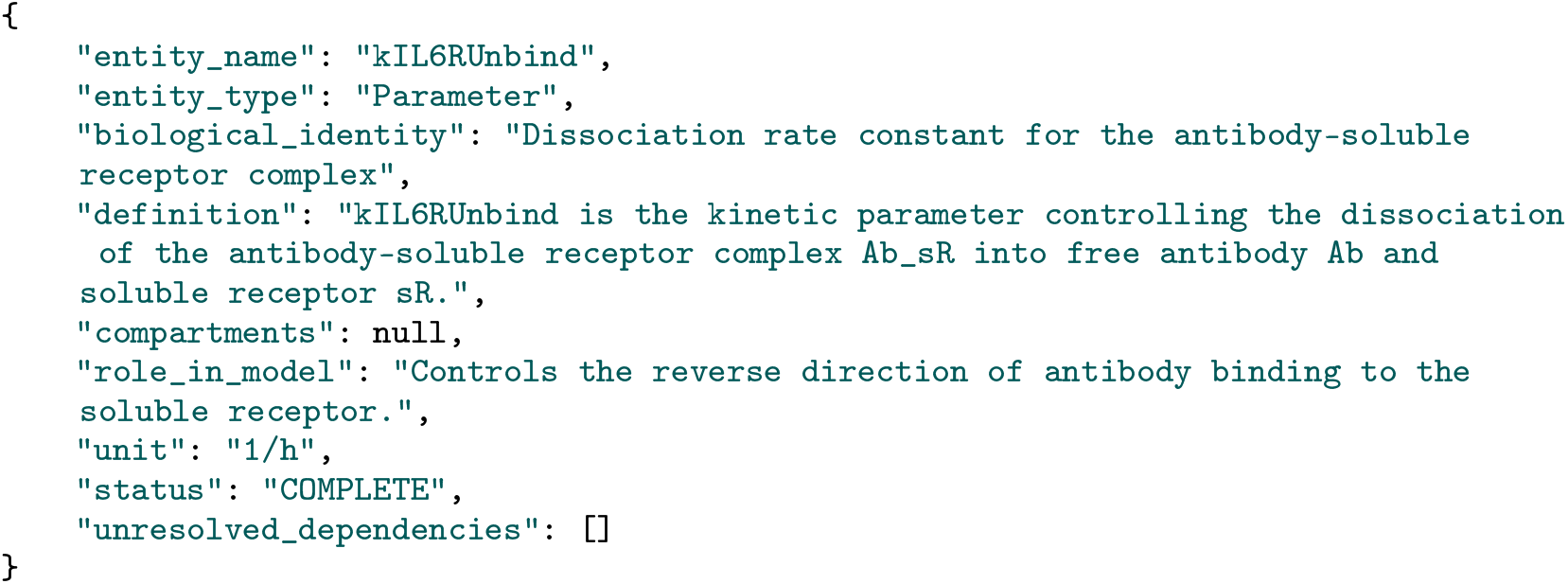

In this representation, the previously unresolved symbols `Ab{serum}`, `sR{serum}`, and `Ab_sR{serum}` have been explicitly interpreted as the free antibody, soluble receptor, and antibody– soluble receptor complex, respectively. The definition therefore no longer depends on unexplained model symbols, and the entity is assigned the status `COMPLETE`. Accordingly, the `unresolved_dependencies` field is empty.

### A.4 The Entity Dictionary and Minified Index

After the multipass refinement stage is complete, the pipeline produces the Entity Dictionary, containing one entry for every model entity. Each entry includes the following fields: `entity_name`, `entity_type`, `biological_identity`, `definition`, `compartments` (filled only for species), `role_in_model`, `status`, `unresolved_dependencies`, `refinement_pass`, `unit`, and `initial_value`.

The following example shows the complete dictionary entry generated for the parameter `kIL6RUnbind` after the refinement process has converged:

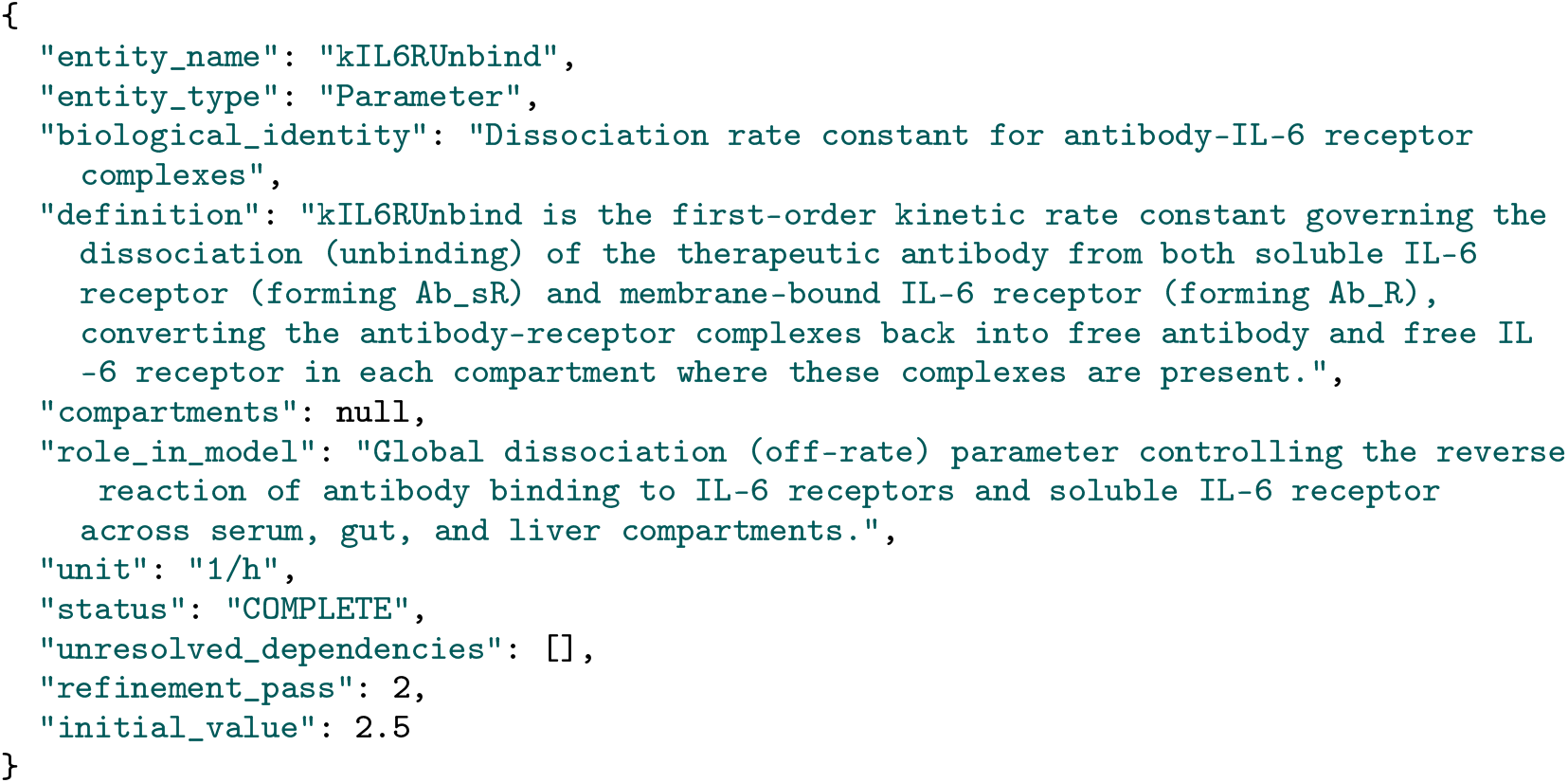

Compared with the intermediate representations in Appendix A.3, this final dictionary entry contains a fully resolved semantic description of the model entity.

The Minified Index is derived from the Entity Dictionary. The two artifacts are best compared on a single entity, so the remainder of this section uses the dosing parameter `DoseQ2W`. Its Entity Dictionary entry is:

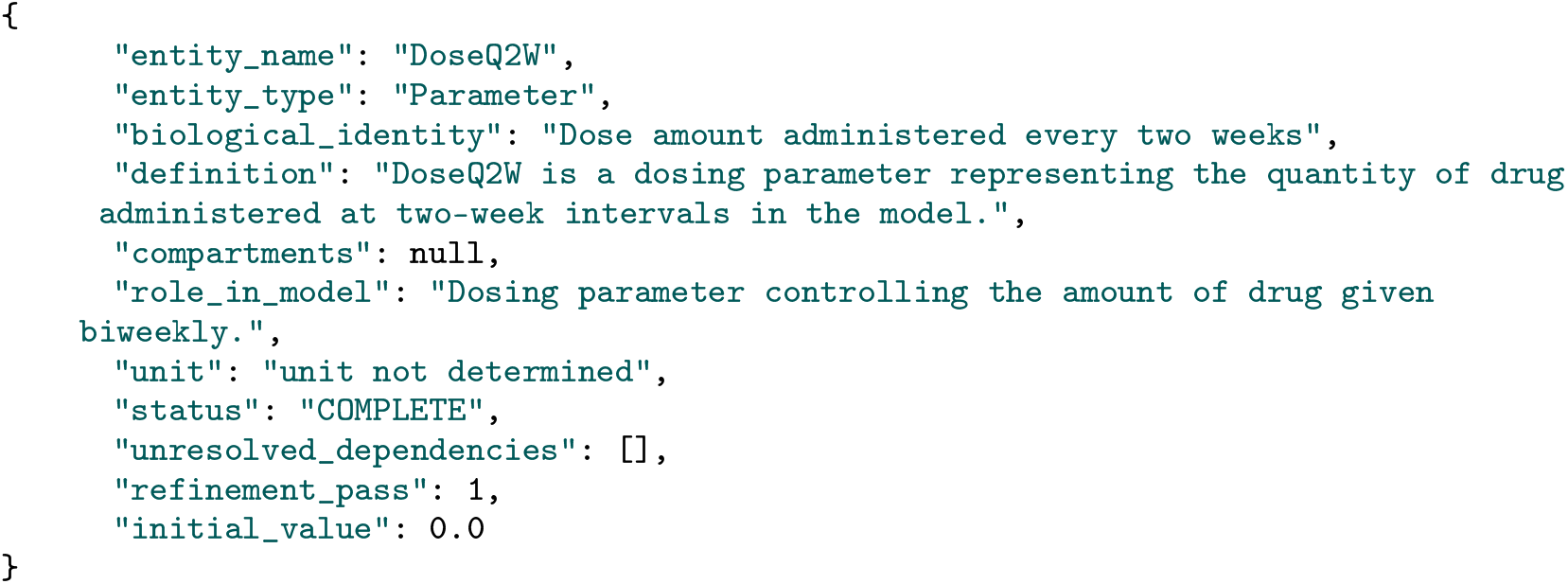

The Minified Index stores a compressed representation of the same entity by retaining only the information required for semantic retrieval and entity matching. The corresponding entry is:

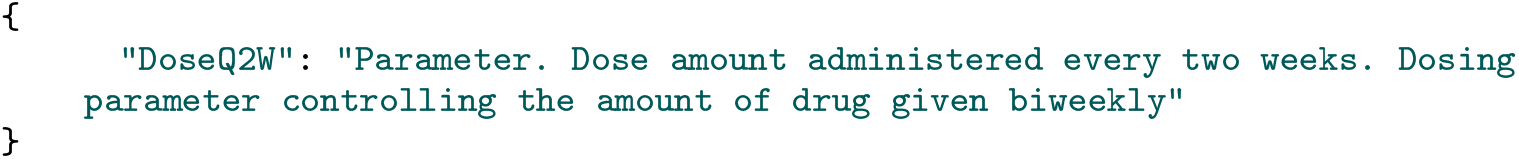

Compared with the Entity Dictionary, the Minified Index omits information such as numerical values, refinement history, unresolved dependencies, and detailed biological definitions. Instead, it retains only the semantic attributes required to identify and match model entities with natural-language descriptions extracted from the literature. This compact representation enables downstream agents to efficiently search across all model entities before retrieving the corresponding Entity Dictionary entry when additional context is required.

### A.5 Example of an Extracted Scenario

The following JSON object illustrates how a scenario is stored together with its source evidence and relevant information:

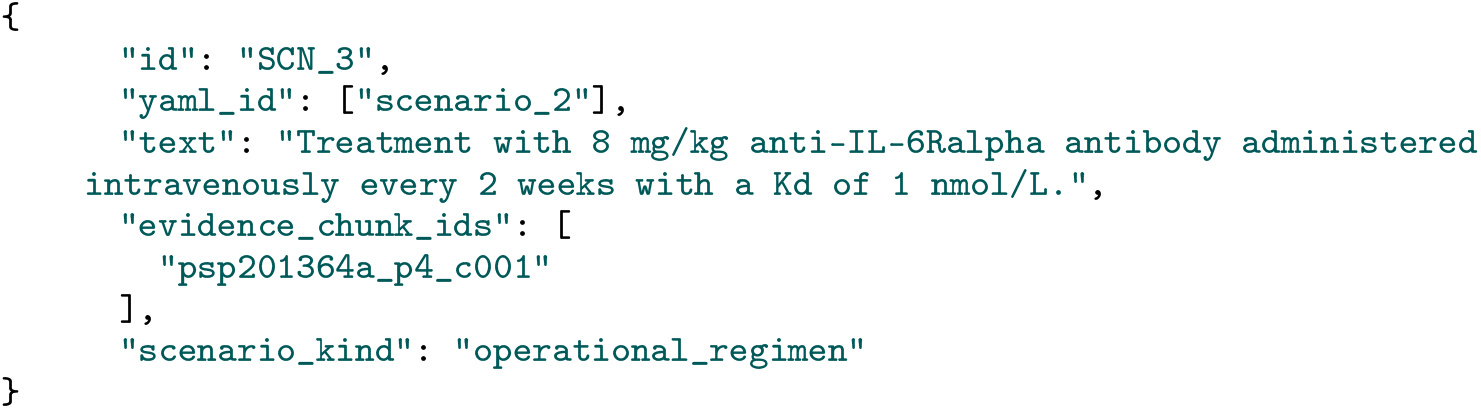

The field `id` provides the internal identifier assigned by the extractor, while `yaml_id` links the extracted scenario to the corresponding scenario identifier in the source model configuration, when available. The `text` field preserves the natural-language scenario extracted from the paper. The `evidence_chunk_ids` field records the exact text chunk from which the scenario was derived, enabling traceability back to the source publication. Finally, `scenario_kind` classifies the scenario according to its functional role; in this example, `operational_regimen` indicates that the scenario describes an executable treatment regimen.

### A.6 End-to-End Example: Mapping an Extracted Scenario into Simulation-Ready Parameters

The following example illustrates the complete mapping process performed by the mapper. The goal of this stage is not to summarize the scenario or explain its biology. Instead, it converts an already extracted scenario into a set of concrete model parameters that can be directly applied to the QSP model.

Consider the following extracted scenario, in the form produced by the Scenario Extractor (Appendix A.5):

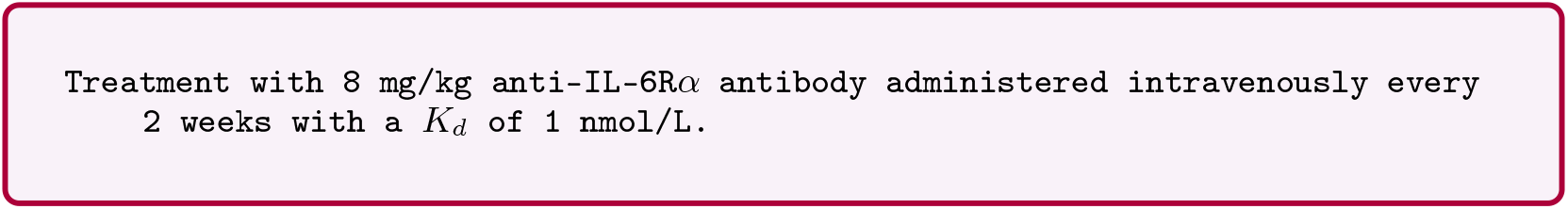

Although this scenario is perfectly understandable to a human reader, it is not directly executable by the model. The model requires exact parameter names and numerical values rather than natural language. The mapper therefore analyzes each component of the scenario independently and translates it into the corresponding model representation.

The Orchestrator first decomposes the scenario into independent work orders, one for each entity that must be resolved:

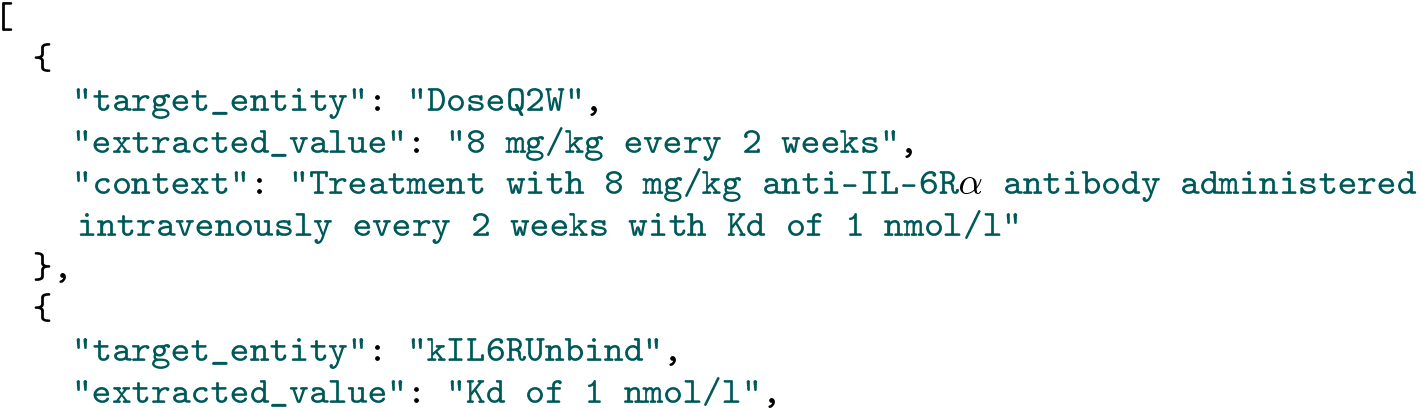
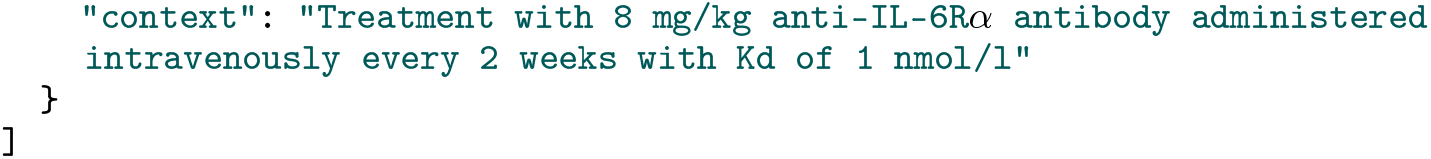

For the first work order, the scenario specifies `every 2 weeks` dosing. The model contains two dosing parameters, `Dose` and `DoseQ2W`, where the second is reserved for the every-two-weeks regimen. The Orchestrator agent therefore targets `DoseQ2W`.

For the second work order, the scenario specifies *K*_*d*_ = 1 nmol/L. The QSP model contains no parameter named *K*_*d*_. Receptor binding is instead represented by two kinetic parameters, `kIL6RBind` and `kIL6RUnbind`, which hold the association and dissociation rate constants of that reaction. The Orchestrator therefore targets `kIL6RUnbind`, the dissociation rate, as the entity the reported affinity constrains.

The Solver then resolves the two work orders in turn and assembles the result.

#### Step 1: Convert the administered dose

The scenario specifies the administered dose as `8 mg/kg`. A **Dictionary Lookup** on `DoseQ2W` shows that the parameter expects the total amount of drug administered rather than a weight-normalized dose, and that no unit is recorded for it in the model. An **Index Lookup** for a body-weight entity returns no match, so the Solver raises two **Human Augmentation Interrupts**, one for the body weight and one for the missing unit, each carrying a default proposal inferred from the context: 70 kg and milligrams. After the expert confirms both, the **Python Calculation Tool** computes the corresponding total dose:

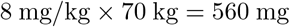

which becomes the model parameter (`DoseQ2W = 560`). At this point, one piece of the natural-language scenario has been transformed into an exact numerical model input.

#### Step 2: Convert the affinity constraint

The scenario reports the affinity as *K*_*d*_ = 1 nmol/L. A **Dictionary Lookup** on `kIL6RUnbind` returns its full definition as the dissociation rate constant, an **Index Lookup** for the corresponding association rate constant returns `kIL6RBind`, and a second **Dictionary Lookup** returns that parameter’s definition and value. Together these fix the relationship 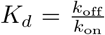. The model already contains `kIL6RBind = 1000`. Using the extracted scenario value *K*_*d*_ = 1, the **Python Calculation Tool** computes *k*_off_ = *K*_*d*_ *× k*_on_ = 1 *×* 1000 = 1000.

Therefore the simulation-ready parameter becomes `kIL6RUnbind = 1000`. The scenario never names this parameter. The Orchestrator agent selected it as the entity the reported affinity constrains, and the Solver agent derived its value from the association rate constant already present in the model.

#### Step 3: Assemble the simulation-ready configuration

After processing every part of the scenario, the mapper combines all parameter updates into the final simulation configuration:

~~~
parameters:
  DoseQ2W:
     value: 560
     unit: mg
  kIL6RUnbind:
     value: 1000
     unit: 1/h
~~~

This configuration can now be passed directly to the QSP model without any additional interpretation.

This example illustrates the complete reasoning process performed by the mapper. Rather than treating the scenario as one block of text, it decomposes it into individual concepts and maps each concept independently.

### A.7 Talk2Biomodels Run Logs

This section reports the full run logs summarized in Table 1. We regenerate two scenarios with Talk2Biomodels in place of the Scenario Mapper, once with GPT-4.1-mini and once with o3 as the backing model, ten runs each. The first scenario is the BIOMD0000000537 every-two-weeks arm, “Treatment with 8 mg/kg anti-IL-6R*α* antibody administered intravenously every 2 weeks with *K*_*d*_ of 1 nmol/L”, for which the correct configuration is `kIL6RUnbind` = 1000 and `DoseQ2W` = 560.

Each run is scored on the first call the agent makes to the `BASICO` simulator: we report the variables and values it submits for execution. The Scenario Mapper is scored on the same basis.

With GPT-4.1-mini as the backing model, no run performs the required unit conversion, and runs variously select the wrong parameter, omit a required one, or submit a miscalculated value, despite the agent having tools available for retrieving model information and variables. Where a call errors, the agent does retry, dropping or substituting the offending variable, but no run reaches the correct configuration under either scoring rule. Every run therefore executes successfully while remaining silently wrong:

**Table 2.** Talk2Biomodels with GPT-4.1-mini on the BIOMD0000000537 every-two-weeks scenario. Values are those submitted to `BASICO` on the first simulator call; a blank cell means the parameter was not submitted. The correct configuration is `kIL6RUnbind` = 1000 and `DoseQ2W` = 560.

| Run | Run Status | DoseQ2W | Dose | Ab |
| --- | --- | --- | --- | --- |
| 1 | Error: kd parameter not found |  |  |  |
| 2 | ok | 8 |  |  |
| 3 | ok |  |  | 8 |
| 4 | ok | 1 | 8 |  |
| 5 | ok |  |  | 8 |
| 6 | Error: kd parameter not found |  |  |  |
| 7 | Error: “anti_IL6R” not found |  |  |  |
| 8 | Error: “anti_IL6R” not found |  |  |  |
| 9 | Error: kd parameter not found | 8 |  |  |
| 10 | $k_{IL6RBind}$: 1, $k_{IL6RUnbind}$: 0.001 | 8 | 0 | |

The same scenario with o3 as the backing model. `kIL6RUnbind` is identified correctly in 9 of 10 runs, but the dose is not: its unit is not determined in the model, and the retrieved context does not disambiguate `Dose` from `DoseQ2W`. Four runs reach the correct configuration; two set both dosing routes, one of them also repeating the schedule as every 14 hours; three submit a dose derived under a different unit assumption (4000, 480, 1296); one submits no change. Every run executes without error:

**Table 3.**
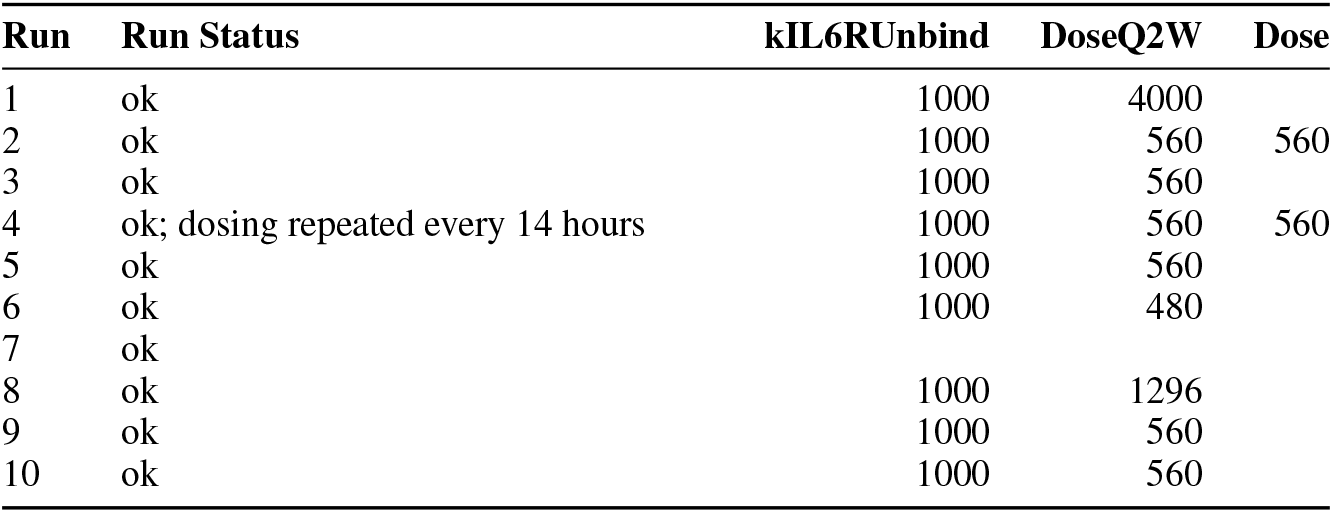
Talk2Biomodels with o3 on the same scenario. Values are those submitted to `BASICO` on the first simulator call; a blank cell means the parameter was not submitted. The correct configuration is `kIL6RUnbind` = 1000 and `DoseQ2W` = 560.

The second scenario uses BIOMD0000000913, a model whose entity names are not self-explanatory.

The scenario is “Treatment with a stabilized combination chemotherapy formulation that exhibits half the baseline degradation rate”, and the correct configuration is `gamma` = 0.45, half its curated initial value of 0.9. We run it first with o3 as the backing model.

`gamma` is the first-order clearance rate constant of the chemotherapy drug. That meaning is not recoverable from the name; it follows from the reaction in which the parameter appears.

Each value o3 submitted is exactly half the curated initial value of the parameter it selected (`d` 5, `s` 0.3, `u` 3, `beta` 0.012, `w` 0.0002): the instruction and the arithmetic are recovered in every run, and only the choice of entity fails.

**Table 4.**
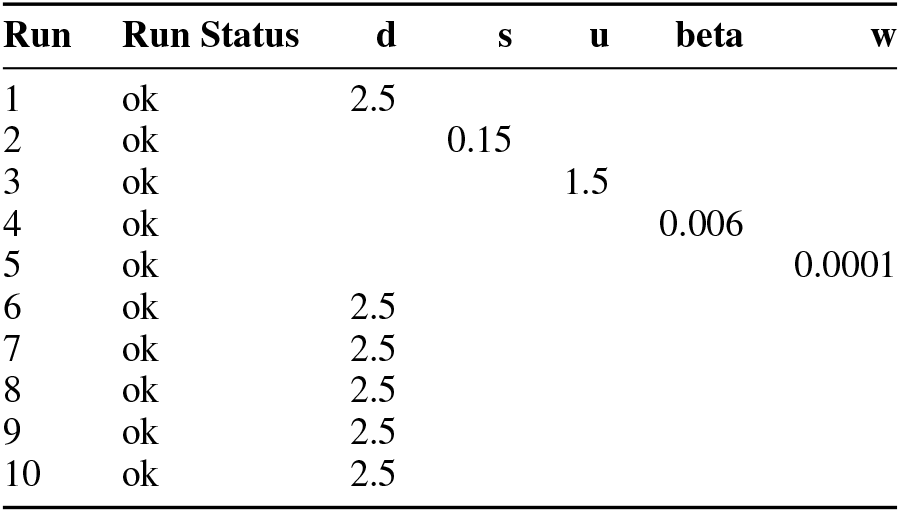
Talk2Biomodels with o3 on the BIOMD0000000913 scenario. Values are those submitted to `BASICO` on the first simulator call; a blank cell means the parameter was not submitted. The correct configuration is `gamma` = 0.45

None of the parameters selected instead governs drug clearance: `d` is the maximum tumor clearance rate, `s` the immune efficacy half-saturation constant, `u` the density-dependent CTL death rate, `beta` the natural lymphocyte clearance rate, and `w` the IL-2-stimulated IL-2 production rate.

We then repeat the same scenario on BIOMD0000000913 with GPT-4.1-mini as the backing model.

Unlike the reasoning model, the smaller model was entirely consistent across runs: it selected `d` and assigned it the same value in every run. It never reached `gamma`. Consistency across runs is therefore not sufficient on its own; the same configuration was reproduced ten times and was wrong ten times.

**Table 5.** Talk2Biomodels with GPT-4.1-mini on the BIOMD0000000913 scenario. Values are those submitted to `BASICO` on the first simulator call. The correct configuration is `gamma` = 0.45

| Run | Run Status | d |
| --- | --- | --- |
| 1 | ok | 2.5 |
| 2 | ok | 2.5 |
| 3 | ok | 2.5 |
| 4 | ok | 2.5 |
| 5 | ok | 2.5 |
| 6 | ok | 2.5 |
| 7 | ok | 2.5 |
| 8 | ok | 2.5 |
| 9 | ok | 2.5 |
| 10 | ok | 2.5 |

Both backing models converge on the same substitution, and both do so silently: all twenty runs on this model execute without error, none reports a failure to identify the target, and in each the submitted value is exactly half the curated initial value of the parameter selected. The gap is therefore not arithmetic but grounding, which the Entity Dictionary supplies to the Scenario Mapper.

## Notes

### Summary of Updates

Substantial revision superseding all previous versions. Title changed to AXIS: Agentic Extraction and Integration of Simulation Scenarios for Quantitative Systems Pharmacology. Authors added and affiliations updated. Abstract, Introduction, Methods and Discussion rewritten. The Entity Definer is renamed the Semantic Profiler. Results now present an end to end reproduction of a published figure from BioModels 537, and a repeated-run comparison with Talk2Biomodels on BioModels 537 and 913. Figures revised and new appendices added.

